# Hemocyte Viability Assay as an Alternative Method for Testing Bacterial Pathogenicity in Bivalves

**DOI:** 10.1101/2025.11.04.686564

**Authors:** Jaypee S. Samson, David C. Rowley, Marta Gomez-Chiarri

## Abstract

Bacterial pathogens cause disease outbreaks in hatcheries, leading to larval mortality and significant economic losses. One of the major challenges in studying bacterial pathogenesis in larvae is the limited availability of healthy larvae throughout the year. This seasonal constraint delays testing and impedes progress in understanding bacterial infections, disease management, and the development of effective treatments for bivalve aquaculture. To address this challenge, a hemocyte viability assay was developed as an alternative for testing bacterial pathogenesis. A resazurin-based assay was optimized using eastern oyster hemocytes, and lethal concentration (LC)_50_ values were determined for the bacterial pathogen *Vibrio coralliilyticus* RE22 (121 ± 2 multiplicity of infection, MOI), probiont *Phaeobacter inhibens* S4 (2,157 ± 56 MOI), and several bacterial isolates from shellfish hatcheries. The most virulent isolates had LC_50_ values ranging from 53 to 195 MOI. To validate the assay, the isolates were tested on oyster larvae, confirming that bacterial isolates with LC_50_ <200 MOI against hemocytes also caused larval mortality, while isolates exhibiting LC_50_ values ≥690 MOI showed no adverse effects on oyster larvae. This high-throughput hemocyte viability assay provides a rapid and reliable alternative to larval assays, particularly during the off-season for hatcheries, facilitating year-round research on bacterial pathogenicity in bivalves.

## INTRODUCTION

Bivalves are important in fisheries, aquaculture, and ecosystem structure and function (Fernández Robledo et al. 2019; Willer et al. 2021; Mesquita et al. 2023). As filter feeders, bivalves acquire a rich and diverse bacterial commensal microbiota including species of *Vibrio, Pseudomonas, Acinetobacter, Photobacterium, Moraxella, Aeromonas, Micrococcus* and *Bacillus* (Kueh and Chan 1985; Zannella et al. 2017). Many of these bacterial species play a beneficial role to the bivalve host, contributing to the digestion of food, including the breakdown of complex carbohydrates and the production of essential amino acids, and the detoxifying of pollutants like heavy metals and organochlorines (Masanja et al. 2023). A few of those bacterial species, however, are pathogenic to bivalves, constituting a leading cause of disease outbreaks in bivalve larviculture systems and resulting in significant mortality and economic losses (Zannella et al. 2017). These bacteria can rapidly proliferate in aquaculture environments, often exploiting stress or environmental fluctuations to infect and overwhelm bivalve hosts (Elston et al. 2008; Dubert et al. 2017). Pathogenic *Vibrio* spp. are particularly well-characterized, producing virulence factors like hemolysins, proteases, and ciliostatic toxins causing deciliation, loss of velar epithelium, and abnormal swimming behavior (Tubiash et al. 1965; Dubert et al. 2017; Gómez-Chiarri et al. 2025).

Experimental larval challenges are commonly used to identify and characterize bacterial pathogens of bivalve larvae (Estes et al. 2004; Gómez-León et al. 2008; Rojas et al. 2019; Wang et al. 2021). The time-consuming nature of these assays, combined with the limited availability of healthy bivalve larvae throughout the year, presents a major challenge in studying the pathogenesis of bacterial isolates, their effects on the larvae, and the development of effective management strategies for bivalve aquaculture. The seasonality of larval production is further complicated by environmental fluctuations and disease pressures that can eradicate or depress larval cohorts (Elston et al. 2008; Dubert et al. 2017; Gradoville et al. 2018), leading to gaps in supply and challenges for experimental replication and validation. Alternative high-throughput methods that are more consistent, accessible, repeatable, and faster are essential. Assays based on measuring the effect of bacterial pathogens on adult bivalve hemocyte viability may provide such an alternative.

Bivalves have an open circulatory system in which hemolymph, containing hemocytes, bathes all the organs before returning to the heart via sinuses. These circulating hemocytes are primarily responsible for defending against pathogens through mechanisms such as phagocytosis and encapsulation (Fisher 1986; Canesi et al. 2002; Gerdol et al. 2018). Additionally, antibacterial effectors, opsonins, nonspecific hydrolytic enzymes, and toxic oxygen intermediates found in bivalve hemolymph coordinate the immune response (Song et al. 2010; Gerdol et al. 2018; Chong 2022). Hemocytes can recognize, locate, ingest, transport, and digest foreign particles, playing a central role in phagocytosis, encapsulation, inflammation, and wound repair (Fisher 1986; Gerdol et al. 2018). Hemocyte viability is a measure of the proportion of alive cells to evaluate overall hemocyte health. Viability assays can be assessed based on cellular metabolism, enzyme activity, or cell membrane integrity (Kamiloglu et al. 2020). Double fluorescence staining with membrane-permeable nuclear dyes, such as SYBR, and membrane-impermeable dyes, such as propidium iodide (PI), coupled with fluorescence microscopy or flow cytometry, has been commonly used to determine hemocyte viability, but these assays are not quantitative or require complex instrumentation (Allam et al. 2001). Alternatively, the Alamar Blue assay, also known as the resazurin assay, offers an attractive means of assessing hemocyte viability. This assay relies on the reduction of resazurin to resorufin, a fluorescent compound that serves as an indicator of cellular metabolic activity (O’Brien et al. 2000). Resazurin-based assays have been widely used to measure cell viability and cytotoxicity in a range of biological and environmental systems and several cell types, including bacteria, yeast, fungi, protozoa, and cultured mammalian and piscine cells (O’Brien et al. 2000; Rampersad 2012; Petiti et al. 2025).

The goal of this work was to develop a high-throughput hemocyte viability assay for assessing bacterial virulence in bivalves without relying on the seasonal availability of bivalve larvae. The assay workflow involves exposure of adult bivalve hemocytes to serial dilutions of bacteria allowing for bacterial-hemocyte interaction, followed by treatment with antibiotics to eliminate bacterial metabolic activity, and, finally, measurement of hemocyte cellular metabolic activity via reduction of resazurin dye. The use of this assay as a screening tool for pathogens of bivalve larvae was validated using larval bacterial challenge assays. This rapid, high-throughput, plate-based resazurin assay will facilitate the identification and characterization of bivalve bacterial pathogens.

## MATERIALS AND METHODS

### Bacteria Isolation and Identification

Bacteria were isolated from larvae of the eastern oyster *Crassostrea virginica* and northern quahog *Mercenaria mercenaria* from four hatcheries located along the Atlantic Coast of the United States. The larvae were rinsed and incubated in filtered sterile seawater (FSSW, 3% sea salts, Instant Ocean) overnight at room temperature (20 to 22°C) with aeration before bacterial isolation. After overnight incubation, swimming larvae were filtered through a 40-µm mesh and washed with FSSW. A 100 µL aliquot of the concentrated larvae was combined with 100 µL FSSW and macerated using a sterile grinder or pipette tip until the sample became homogeneous. A 100 µL aliquot of the larvae homogenate was then taken for serial dilution. Then, 20 µL of each serial dilution was plated onto various media, including a marine yeast peptone (YP30; 0.5 % yeast extract, 0.1% peptone, 3% sea salts, 1.5 % agar; pH 7.6) (Karim et al., 2013), marine agar 2216 (BD Difco), seawater tryptone agar (SWT, 0.5% tryptone, 0.3% yeast extract, 0.3% glycerol, 3% sea salts, 1.5% agar; pH 7.6), and nutrient-free agar plates (3% sea salts, 1.5% agar; pH 7.6), and incubated for 2-5 days at 27°C. Morphologically distinct colonies were selected and re-streaked onto sterile agar plates to ensure the isolation of pure strains.

Bacterial DNA was extracted following a modified protocol (Li et al. 2023). Briefly, an overnight culture (500 µL) of each isolate was harvested and resuspended in DNase-free water with 0.1 g of 0.1 mm glass beads (Sigma-Aldrich, USA). The cells were heated at 80°C at 500 rpm for 10 min, followed by RNase treatment (Qiagen, Germany). Bacterial isolates were identified by 16S rRNA sequencing using universal primers 27F (5’-AGAGTTTGATCCTGGCTCAG-3’) and 1492R (5’-GGTTACCTTGTTACGACTT-3’) (Weisburg et al. 1991). PCR reactions were prepared in a total volume of 25 µL, comprising 5 µL of extracted DNA, 12.5 µL of PCR mastermix (Thermo Scientific), 1 µL each of forward (27F) and reverse (1492R) primers, and 5.5 µL of nuclease-free water. Amplification was performed using the following thermal cycling conditions: initial denaturation at 95°C for 3 min; 30 cycles of denaturation at 95°C for 30 seconds, annealing at 55°C for 30 seconds, and extension at 72°C for 90 seconds; followed by a final extension at 72°C for 5 min. PCR products were purified using a Qiagen PCR purification kit following the manufacturer’s protocol and sequenced using a Applied Biosystems 3500xL Genetic Analyzer (Thermo Fisher Scientific, USA) at the RI-INBRE Centralized Research Core Facility (University of Rhode Island, Kingston, RI). Sequence identification was conducted by comparing obtained sequences to the NCBI 16S reference database using BLAST (Camacho et al. 2009).

### Overview of the Hemocyte Viability Assay

The workflow for the hemocyte viability assay is illustrated in Fig. 1. The assay relies on incubating hemocytes isolated from adult eastern oysters (Fig. 1A) with dilutions of the bacterial isolates to allow for bacterial – hemocyte interaction (Fig. 1B). After this period of incubation, the hemocyte – bacterial mix is incubated with antibiotics to kill the bacteria (Fig. 1C). Finally, a resazurin-based dye (CellTiter-Blue, Promega) is added to evaluate hemocyte viability (Fig. 1C).

**Figure 1.**
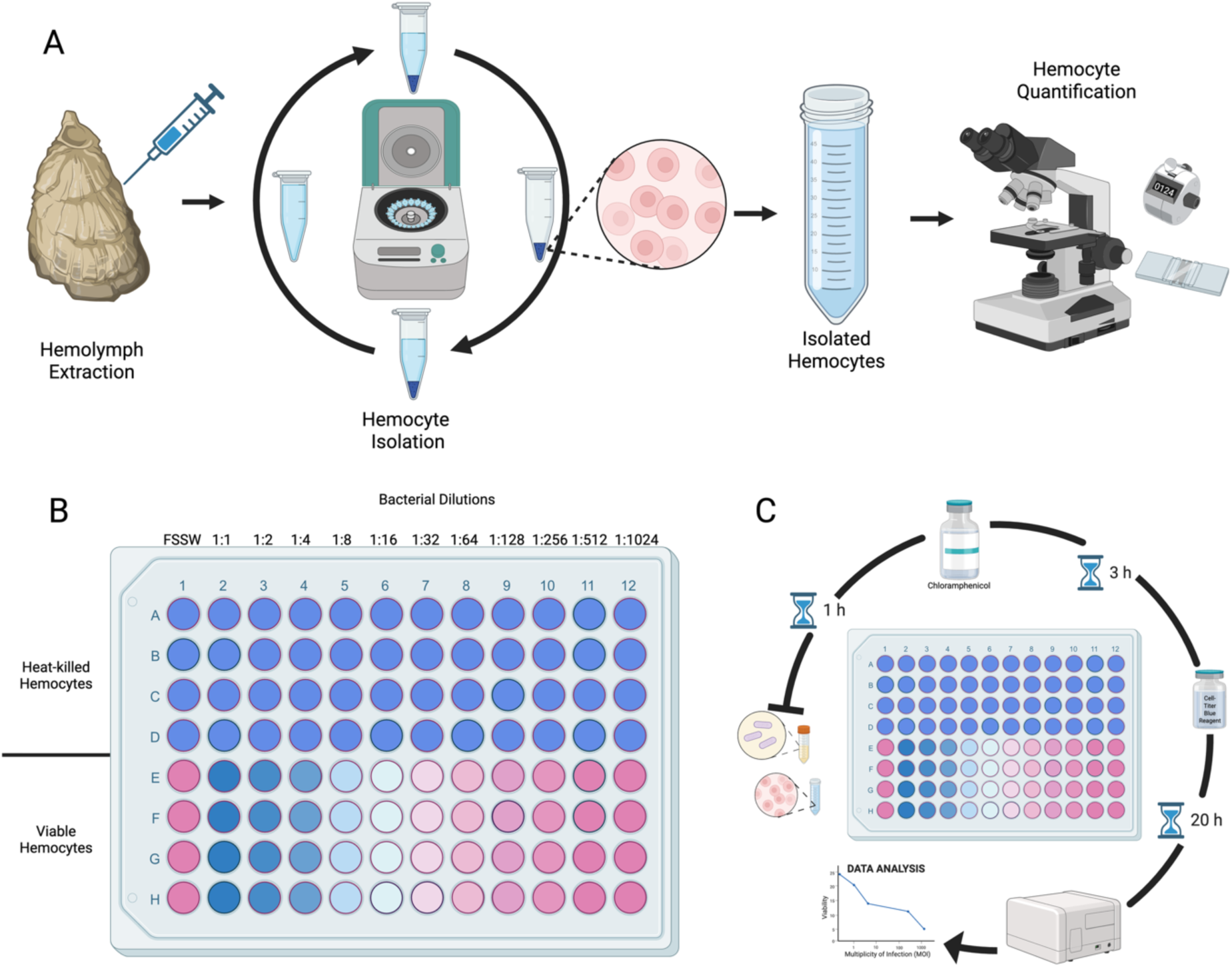
Overview of the hemocyte viability assay including the (A) isolation, preparation, and quantification of cell concentration; (B) plate layout, with half of the plate for the incubation of live hemocytes with bacterial dilutions, and half of the plate for heat killed hemocytes plus bacterial dilution controls; and (C) the procedure of the hemocyte viability assay using the resazurin dye. Figure created with BioRender.com.

### Preparation of Bacterial Dilutions

Bacterial isolates (Table 1) were cultured overnight on YP30 agar plates at 27°C for 18-24 hours. After incubation, individual colonies were picked and transferred to liquid YP30 medium, which was incubated at 27°C with shaking (150 rpm) for 18-24 hours. Following the incubation, the bacterial cultures were centrifuged at 16,000 ′ g for 5 min to pellet the cells. The supernatant was discarded, and the bacterial pellet was washed twice with FSSW. The cell pellet was then resuspended in FSSW, and the bacterial concentration of the stock was adjusted to 108 CFU mL-1 based on optical density (OD_600_). Two-fold bacterial dilutions were prepared in sterile 96-well plates. The final bacterial concentration in each well was verified by spot-plating appropriate dilutions onto YP30 agar plates and determining colony counts (Karim et al. 2013).

**Table 1.** List of bacterial isolates used in the study.

| ISOLATE TESTED | ISOLATION SOURCE | REFERENCE |
| --- | --- | --- |
| *Algoriphagus yeomjeoni* DEN5 | *C. virginica* | This study |
| *Cellulophaga lytica* CH30 | *C. virginica* | This study |
| *Glutamicibacter soli* CLAM16 | *M. mercenaria* | This study |
| *Marinomonas gallaica* CLAM9 | *M. mercenaria* | This study |
| *Phaeobacter inhibens* S4 | *C. virginica* | Karim et al., 2013 |
| *Pseudooceanicola nitratireducens* NEH7 | *C. virginica* | This study |
| *Tenacibaculum ascidiaceicola* CLAM15 | *M. mercenaria* | This study |
| *Vibrio alginolyticus* DEN30 | *C. virginica* | This study |
| *Vibrio chagasii* CH4 | *C. virginica* | This study |
| *Vibrio chagasii* CLAM14 | *M. mercenaria* | This study |
| *Vibrio coralliilyticus* RE22 | *C. gigas* | Estes et al., 2004 |
| *Vibrio cyclitrophicus* CLAM19 | *M. mercenaria* | This study |
| *Vibrio fortis* CH6 | *C. virginica* | This study |
| *Vibrio neptunius* DEN11 | *C. virginica* | This study |
| *Vibrio rotiferianus* CH3 | *C. virginica* | This study |
| *Vibrio tasmaniensis* CLAM18 | *M. mercenaria* | This study |
| *Vibrio toranzoniae* CH7 | *C. virginica* | This study |
| *Vibrio tubiashii* DEN41 | *C. virginica* | This study |
| *Vibrio tubiashii* DEN12 | *C. virginica* | This study |

### Preparation of Oyster Hemocytes

Hemocyte isolation was performed on adult eastern oysters collected from Rhode Island farms using aseptic conditions (Fig. 1A). The oyster shells were scrubbed with a sterilized soft-bristle brush to remove surface biofilm, rinsed in freshwater and sprayed with 70% ethanol. A small notch was made on the edge of the shell at the height of the adductor muscle using flame-sterilized pliers. Next, a sterile 1.5" 21G needle attached to a 3 mL syringe was primed with 100 µL of ice-cold FSSW, ensuring that air bubbles were removed from the syringe. The primed needle was inserted into the adductor muscle notch, and approximately 1 mL of hemolymph was extracted from each oyster. The needle was rotated gently during extraction to help remove tissue adhering to the needle. To remove tissue debris, hemolymph samples were passed through a 40 µm mesh filter into a 50 mL sterile Falcon tube kept on ice. The pooled hemolymph of 6-12 oysters was then centrifuged at 600 × g for 3 min at 4°C to pellet the hemocytes. The hemocytes were washed twice in FSSW and finally resuspended in FSSW. Hemocyte concentration and viability were evaluated using a hemocytometer, and the concentration was adjusted to 2.0 × 10^6^ cells mL^-1^. For heat-killed hemocyte preparations (blank control), half of the total hemocyte suspension volume was placed in a water bath at 90°C for 10 min. Live and heat-killed (HK) hemocyte solutions were kept on ice until used in the assay.

### Hemocyte Viability Assay Development

The assay was optimized to determine the appropriate hemocyte and antibiotic concentrations, as well as the incubation times for each phase: hemocyte–bacteria interaction, antibiotic treatment, and resazurin metabolism. The final test conditions for the optimized assay were selected based on the most optimal signal to noise ratio and lowest coefficient of variation. The optimized protocol is as follows. Using black clear-bottom 96-well plates (Corning, USA), 50 µL of either live or heat-killed hemocytes were added to designated wells of the plate to achieve a final concentration of 1.0 × 10^6^ cells mL^-1^ (Fig.1B). Fifty µL of serial dilutions of the test bacterial suspensions (each in quadruplicate) were added to wells containing hemocytes to achieve a range of multiplicity of infection (MOI). The MOI was calculated by dividing the bacterial concentration (CFU mL^-1^) by the hemocyte concentration (cells mL^-1^). The plates were incubated for 1 hour at room temperature to allow bacterial interaction with the hemocytes, then treated with chloramphenicol (200 µg mL^-1^ final concentration) for an additional three hours to prevent further bacterial metabolism. Next, wells were treated with 10 µL of resazurin reagent (Promega, USA) and the plates were incubated for 20 hours at room temperature. Plates were gently shaken for 10 seconds and fluorescence was measured (Fluorometer model; BioTek, USA) at an excitation wavelength of 560 nm and an emission wavelength of 590 nm (Fig. 1C).

The relative viability of the hemocytes was determined using the following formula:

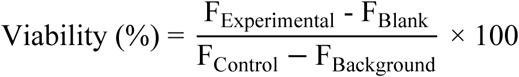

Where:

F_Experimental_= fluorescence readings of wells containing hemocytes with bacterial treatment.
F_Blank_= fluorescence readings of wells containing heat-killed hemocytes with bacterial treatments (accounts for background fluorescence as well as potential residual metabolic activity from bacteria that was not eliminated by the antibiotic treatment).
F_Control_ = fluorescence readings of wells containing hemocytes without bacterial treatment.
F_Background_= fluorescence reading of wells containing heat-killed hemocytes without bacterial treatment.

Linear regression analysis was performed to plot the relationship between MOI and relative viability. Bacterial concentrations (expressed as MOI) resulting in a 50% reduction in hemocyte viability (LC_50_) were determined using the regression equation. All calculations were performed using Microsoft Excel.

### Larval Viability Assays

Larval viability assays were performed using a previously described protocol (Gómez-León et al. 2008) with modifications. Briefly, 2 mL of 6-8-day-old healthy eastern oyster larvae (veligers) were added to 24-well plates at 75-100 larvae mL^-1^. Then, bacterial isolates were added at a final concentration of 10^5^ CFU mL^-1^ and incubated for 16-20 h at room temperature. Larval mortality was determined by visual examination under a microscope (Leica, 400x magnification). Larvae mortality was calculated as follows:

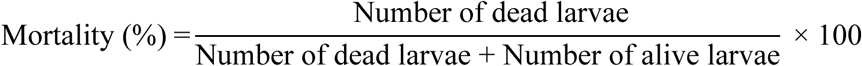

## RESULTS

### Assay Development and Optimization

To establish a reliable and rapid hemocyte viability assay for assessing bacterial virulence, we explored the use of resazurin dye to measure cell survival following treatment with various bacteria. Several parameters were optimized, including hemocyte concentration (ranging from 2.5 × 10^5^ to 2 × 10⁶), bacteria – hemocyte interaction time (1 to 5h), antibiotic treatment (ampicillin, gentamycin, penicillin, chloramphenicol), and resazurin – hemocyte incubation times (from 1 to 24h). Hemocyte concentrations between 1 × 10⁶ and 2 × 10⁶ cells mL^-1^ were determined to yield reliable and reproducible fluorescence measurements (not shown). A concentration of 200 μg mL^-1^ of the broad-spectrum antibiotic chloramphenicol was determined to effectively inhibit the metabolic activity of bacteria without impacting hemocyte viability (Table 2). Fluorescence readings stabilized after 20 hours of incubation of the hemocytes with the resazurin dye, indicating sufficient time for metabolic conversion and accurate viability measurement. Overall, the assay has a 24 h duration, including the bacteria-hemocyte interaction, antibiotic treatment, and incubation with dye (Fig. 1).

**Table 2.** Antibiotic concentration to prevent or reduce bacterial growth.

| Chloramphenicol (ug mL<sup>-1</sup>) | *V. coralliilyticus* RE22 | Hemocyte viability |
| --- | --- | --- |
| 5 | + | + |
| 10 | + | + |
| 20 | + | + |
| 40 | + | + |
| 60 | + | + |
| 80 | + | + |
| 100 | + | + |
| 150 | +/- | + |
| 200 | - | + |
| 300 | - | + |
| 400 | - | + |
| 500 | - | + |

+ = resazurin dye turned into pinkish
- = resazurin dye remained bluish

### Assay Validation Using Known Pathogenic and Probiotic Bacteria

The assay was validated using the larval pathogen *V. coralliilyticus* RE22 and probiont *Phaeobacter inhibens* S4 (Karim et al., 2013). The calculated LC_50_ values, defined as the multiplicity of infection (MOI, ratio of bacteria to hemocytes) required to reduce hemocyte viability by 50%, were determined to be 121 ± 2 for RE22 and 2,157 ± 56 for *P. inhibens* S4 (Fig. 2). The assay was repeated three times, with each bacterial concentration tested in quadruplicate, to calculate inter and intra-assay variability. The coefficient of variation for inter-assay accuracy was determined to be 6.3% and 7.7% and for RE22 and S4, respectively.

**Figure 2.**
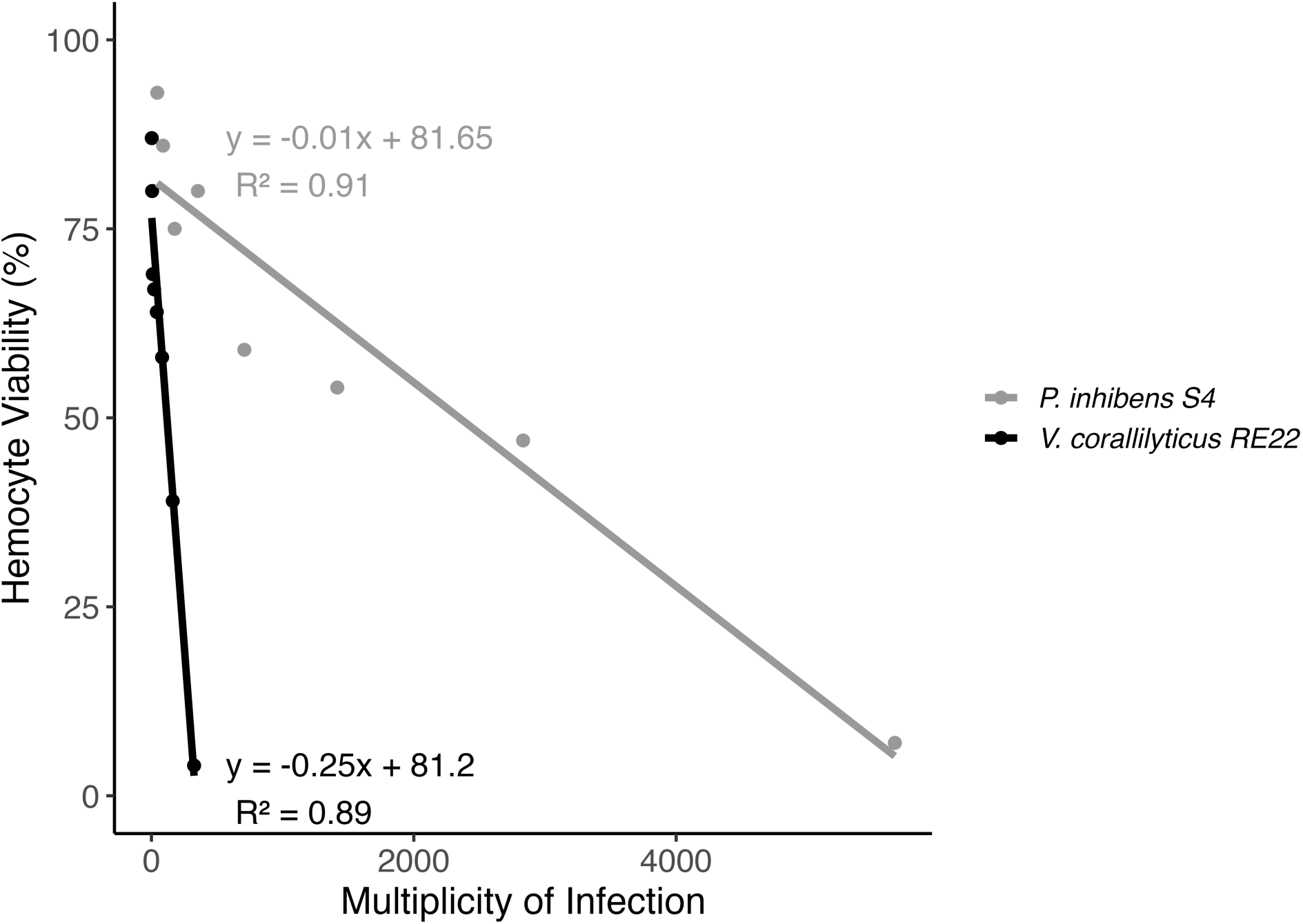
Relationship between multiplicity of infection (MOI) and hemocyte viability measured by resazurin reduction. Linear regression analysis of MOI and hemocyte viability for the pathogen *V. coralliilyticus* RE22 and the probiont *P. inhibens* S4 demonstrates the negative correlation between MOI and hemocyte metabolic activity, indicating reduced hemocyte viability at higher bacterial loads. Data points represent mean values from replicate wells, with fitted regression lines, corresponding equations, and R² values.

### Application to Field Isolates

The assay was further applied to evaluate the virulence of several bacterial isolates isolated from oyster and clam larvae (Table 1). LC_50_ values ranged from 53 to 25,389 MOI (Table 3), indicating a wide spectrum of virulence. Among the most virulent isolates, *V. toranzoniae* CH7 displayed an LC_50_ of 53 MOI, followed by *V. rotiferianus* CH3 (81 MOI), *V. fortis* CH6 (86 MOI), *V. neptunius* DEN11 (129 MOI), *V. chagasii* CH4 (190 MOI), and *C. lytica* CH30 (195 MOI). In contrast, *V. chagasii* CLAM14 and *T. ascidiaceicola* CLAM15 displayed LC_50_ values > 20,000 MOI, suggesting these would be non-pathogenic isolates to eastern oysters.

**Table 3.** Effect of bacterial isolates on adult oyster hemocyte viability and larval mortality. Effect on hemocyte viability is quantified by the concentration of bacteria, as measured by multiplicity of infection (MOI), required to reduce hemocyte viability by 50% (LC_50_). Effect on larvae is measured as the percent mortality experienced in a larval culture after a 20 h incubation with a concentration of 10^5^ CFU ml.

| ISOLATE TESTED | LC<sub>50</sub> (MOI)<br>(Average ± SEM) | Mortality (%)<br>(Average ± SEM) |
| --- | --- | --- |
| *Vibrio fortis* CH6 | 84 | 0.0 ± 0.0 |
| *Vibrio neptunius* DEN11 | 129 | 0.0 ± 0.0 |
| * *Vibrio coralliilyticus* RE22 | 121 ± 2 | 0.9 ± 0.9 |
| *Vibrio chagasii* CH4 | 190 | 18.1 ± 0.1 |
| *Cellulophaga lytica* CH30 | 195 | 23.5 ± 6.5 |
| *Vibrio rotiferianus* CH3 | 81 | 23.9 ± 0.7 |
| *Vibrio toranzoniae* CH7 | 53 | 30.1 ± 0.9 |
| *Vibrio tubiashii* DEN41 | 848 | 45.9 ± 1.7 |
| *Vibrio tubiashii* DEN12 | 4,160 | 46.3 ± 8.1 |
| *Vibrio alginolyticus* DEN30 | 12,204 | 79.7 ± 5.0 |
| *Vibrio cyclitrophicus* CLAM19 | 927 | 93.2 ± 2.2 |
| *Marinomonas gallaica* CLAM9 | 704 | 96.4 ± 1.8 |
| *Pseudooceanicola nitratireducens* NEH7 | 690 | 97.6 ± 0.8 |
| *Vibrio chagasii* CLAM14 | 25,389 | 98.6 ± 1.3 |
| *Tenacibaculum ascidiaceicola* CLAM15 | 20,315 | 99.3 ± 0.7 |
| *Glutamicibacter soli* CLAM16 | 1,913 | 100.0 ± 0.0 |
| * *Phaeobacter inhibens* S4 | 2,157 ± 56 | 100.0 ± 0.0 |
| *Vibrio tasmaniensis* CLAM18 | 877 | 100.0 ± 0.0 |
| *Algoriphagus yeomjeoni* DEN5 | 935 | 100.0 ± 0.0 |

\* Known pathogen and probiotics. SEM: Standard error of the mean.

### Comparison of the Hemocyte Viability Assay with the Larval Assay

To assess the relationship between hemocyte viability and larval mortality, the bacterial isolates were tested against eastern oyster larvae in challenge assays. Larvae were exposed to bacterial isolates for 16 – 20 h at a standardized bacterial concentration of 10^5^ CFU mL^-1^ (Table 2). Oyster larvae challenged with the known pathogen *V. coralliilyticus* RE22 had almost 100% mortality. Similarly, potential pathogenic isolates *V. fortis* CH6 and *V. neptunius* DEN11 also had 100% mortality. In addition, *V. chagasii* CH4, *C. lytica* CH30, *V. rotiferianus* CH3, *V. toranzoniae* CH7, and *V. tubiashii* DEN41 had larval survival below 50%, confirming their pathogenicity in both assays. Conversely, larvae treated with putative commensal or probiotic isolates exhibited survival rates of 96-100%, indicating their safety towards larvae. Bacterial isolates demonstrating LC_50_ values below 200 MOI in the hemocyte assay also exhibited significant pathogenicity towards oyster larvae. The correlation between mortality and LC_50_ of the pathogenic isolates was significant, with a Pearson correlation coefficient of 0.84 (*P* = 0.0021) (Fig. 3). The range of MOI for isolates that did not cause mortality (survival >90%) in the 20 hour larval assay, however, was highly variable, ranging from 690 to 23,389, suggesting that the hemocyte assay is more sensitive than the short term larval assay.

**Figure 3.**
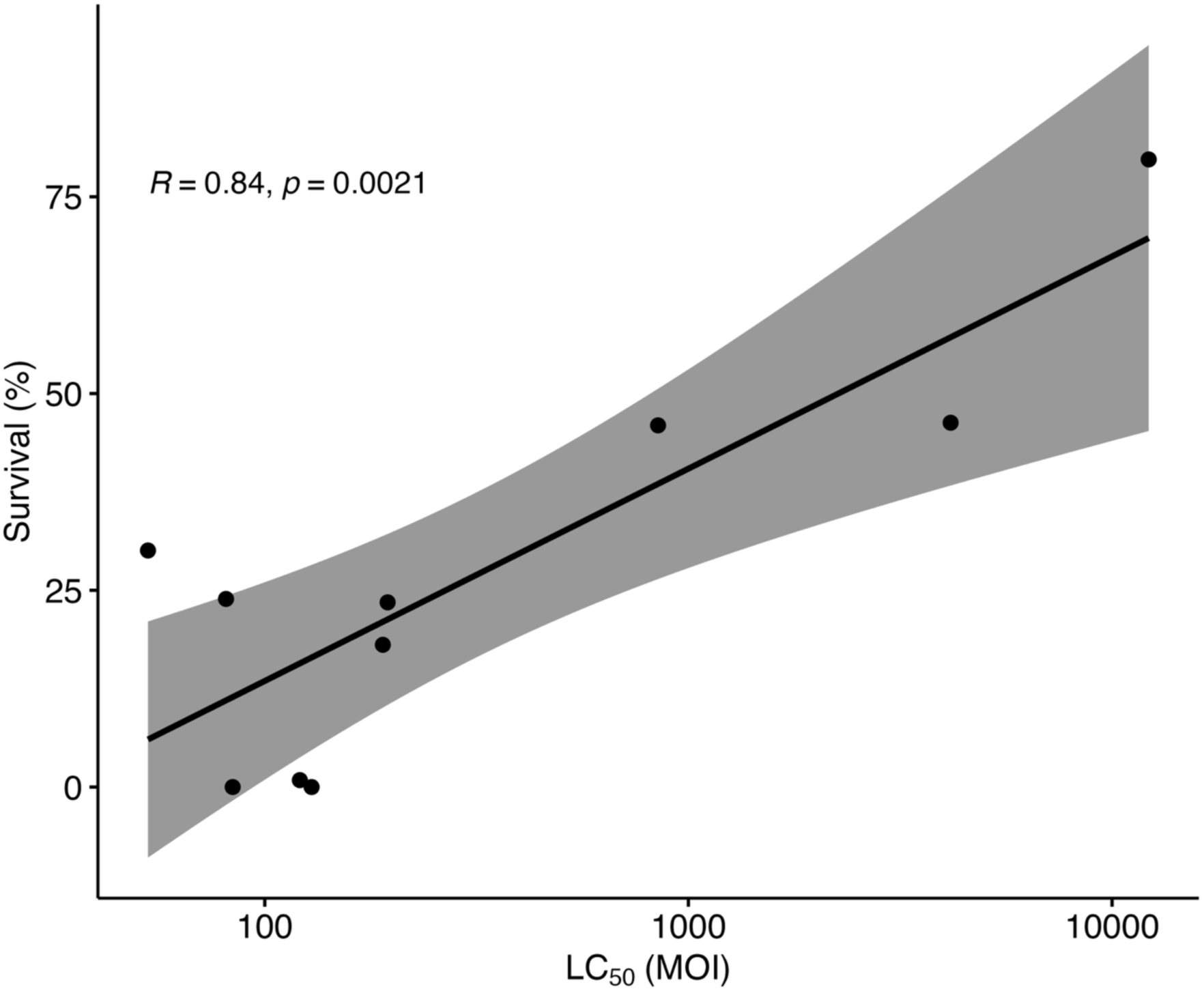
Relationship between hemocyte and larval viability assays. Linear regression of larval survival after treatment with pathogenic isolates (as defined as those causing >20% mortality) and their corresponding LC_50_ values on adult oyster hemocytes based on multiplicity of infection (MOI). The survival percentage is plotted against the LC_50_ values on a logarithmic scale. Larval survival data points represent mean values of 3 wells of larvae treated with 10^5^ CFU mL^-1^ of bacteria for 16-20 hours. Pearson correlation and linear regression with 95% confidence intervals are shown.

## DISCUSSION

This study presents a rapid, high-throughput, and reliable hemocyte viability assay as an alternative to traditional larval assays for assessing bacterial virulence against bivalves. Unlike larval-based methods, which are limited by the seasonal availability of healthy larvae and are time-consuming and variable, this assay provides a consistent and quantifiable measure of bacterial effects on pools of adult hemocytes, which can be accessed year-round. Hence, this assay is an attractive tool to complement traditional larval challenge models for the screening of potential bacterial pathogens and the development of disease management strategies, such as probiotic selection.

To our knowledge, this is the first report of using resazurin-based viability measurement in bivalve hemocytes to screen for potential pathogens and probiotics of larval bivalves and quantify bacterial virulence. By using resazurin reduction as a measure for hemocyte viability, the assay enables sensitive quantification of bacterial virulence in a controlled, reproducible format. One initial challenge encountered during the assay development was interference by bacterial reduction of resazurin. This issue was resolved by the addition of chloramphenicol, a broad-spectrum antibiotic, after allowing for bacterial-hemocyte interaction and bacterial pathogenicity, and the inclusion of the killed hemocyte and bacterial control, which helps identify those bacterial isolates for which the antibiotic treatment failed to eliminate bacterial metabolism. Hemocytes were unaffected by the antibiotic treatment step, similar to previous reports (Lacaze et al. 2015; Matozzo et al. 2016). Hence, the optimized assay is capable of measuring hemocyte reduction of resazurin without interference from the bacteria isolates being evaluated.

### The Hemocyte Viability Assay was Significantly Correlated with the Larval Viability Assay

To validate the hemocyte viability assay, a collection of 19 bacterial isolates from bivalve larvae was tested using both the hemocyte viability assay and the larval viability assay. The two assays showed a significant correlation for those isolates causing mortality in oyster larvae. Isolates *V. toranzoniae* CH7, *V. rotiferianus* CH3, *V. fortis* CH6, *V. neptunius* DEN11, *V. chagasii* CH4, and *C. lytica* CH30, which showed LC_50_ values of <200 MOI, caused ≥ 30% mortality in oyster larvae, confirming the suitability of the hemocyte viability assay to identify larval pathogens. The taxonomic identification of pathogenic isolates using the hemocyte viability assays is consistent with previous reports of pathogenicity toward bivalves (Dubert et al. 2017; Gómez-Chiarri et al. 2025). For example, *V. toranzoniae* and several other *V{Citation}ibrio* spp. (*V. lentus*, *V. crassostreae*, and *V. cyclitrophicus*) caused significantly higher hemocyte cytotoxic rates in the scallop, *Argopecten purpuratus* (González et al. 2025 Apr 1). Similarly, *V. rotiferianus* shows pathogenicity in infection models (Harrison et al. 2022), and *V. fortis* has been implicated in the heat stress-induced mortality of *C. gigas* (Green et al. 2019). Additionally, *V. neptunius* has been identified as an opportunistic pathogen that causes disease in oysters and clams (Galvis et al. 2021), while *V. chagasii* was reported as the causative agent of massive larval mortalities in farmed scallops (*A. purpuratus*) in Chile (Urtubia et al. 2023). Lastly, *C. lytica* has been isolated in the eroded mantle epithelium of juvenile paua (*Haliotis iris*) in New Zealand (Diggles and Oliver 2005).

Similarly, most isolates with LC_50_ values ≥ 690 MOI in the hemocyte viability assay caused no mortality in the 16–20 hour larval assay, indicating overall good concordance between the two assays. Only three of the 19 tested isolates causing mortality in larvae (*V. tubiashii* isolates DEN12 and DEN41 and *V. alginolyticus* DEN30) showed levels of hemocyte viability comparable to isolates that did not cause larval mortality (LC_50_ of 4,160, 848, and 12,204 MOI respectively). This suggests that the pathogenicity of these three isolates to larvae may involve virulence mechanisms not fully captured by hemocyte viability alone, such as tissue-specific interactions or factors expressed only *in vivo*, and/or that adult oyster hemocytes have increased capability for eliminating these strains due to immune system maturation and/or priming (Leonessi et al. 2025). Differences in the LC_50_ in the hemocyte viability assay between the two *V. tubiashii* strains, as well as differences between the two *V. chagasii* strains, is consistent with the well-established principle that *Vibrio* pathogenicity is determined at the strain level, with most species exhibiting a diversity of nonpathogenic and pathogenic strains (Estes et al. 2004; Ushijima et al. 2018; Destoumieux-Garzón et al. 2020; Gómez-Chiarri et al. 2025).

The wide range of values in the hemocyte viability assay for isolates causing no mortality in larvae (LC_50_ from 690 to 25,395 MOI) also suggests that the hemocyte viability assay may be more sensitive than a larval assay of a similar duration (around 24 hours), and that some of the isolates may require longer incubation periods with the larvae to show differential impacts on survival. A higher sensitivity of hemocytes to bacterial pathogenicity compared to larvae may also be due to the additional immune defenses and barriers present in larvae (e.g., mucosal barriers (Fernández Robledo et al. 2019)) and/or differential interactions of these with the more complex larval microbiome (Yeh et al. 2020; Arfken et al. 2021; Masanja et al. 2023; Destoumieux-Garzón et al. 2024; Takyi et al. 2024). Notably, six of the eight bacterial strains causing no mortality in oysters were isolated from clams. Because our hemocyte assays were performed using oyster hemocytes, it is possible that these isolates exhibit host specificity and are pathogenic to clams. Such host-specific interactions are well documented in Vibrio – bivalve systems, where virulence can vary widely depending on the host species, immune cell type, and ecological context (Destoumieux-Garzón et al. 2020; Gómez-Chiarri et al. 2025).

## CONCLUSIONS

This study aimed to develop an alternative assay for assessing bacterial pathogenicity in bivalves, utilizing hemocytes isolated from adult oysters as a substitute for oyster larvae. The results from the hemocyte viability assay developed here correlated well with those from traditional larval challenges, offering a rapid, consistent, and effective tool for testing the pathogenicity of bacterial isolates against bivalves. Some potential limitations of this assay include the fact that this *in vitro* assay may not fully capture the complexity of whole-organism responses in larvae, such as tissue-specific effects or systemic immune responses, a limitation common to most high-throughput *in vitro* screening assays (Ghallab 2013). Additionally, some bacterial species or target host cells may exhibit variable sensitivity to chloramphenicol and/or resazurin (Breijyeh et al. 2020; Vieira-da-Silva and Castanho 2023), potentially affecting assay accuracy. Alternate antibiotics, or even antibiotic cocktails, may be needed in such cases. Despite these limitations, the hemocyte viability assay provides a high-throughput tool for pre-screening isolates for potential pathogenicity and helps solve the bottleneck of larval shortages encountered during the off-season, thereby enabling year-round investigation of potential pathogens and the development of disease mitigation tools for hatcheries. Importantly, this assay can be further adapted to assess the pathogenicity of bacterial isolates in hemocytes from other bivalve species, such as clams, scallops, and mussels. This flexibility makes it a valuable tool for investigating host-specific pathogens across a broader range of bivalve species.

## Acknowledgements

We are grateful to all the research and commercial hatcheries involved in the Bivalve Hatchery Health Consortium, current and previous members of the Gomez-Chiarri lab for their assistance during this study, and all the members of the Probiotics Working Group at the University of Rhode Island.

